# BAP1 loss impairs Non-Homologous End Joining DNA repair promoting genomic instability

**DOI:** 10.64898/2026.02.17.706330

**Authors:** Sabrina Caporali, Carla Schuy, Alessio Butera, Melanie Rall-Scharpf, Nils Geese, Lisa Wiesmüller, Ivano Amelio

## Abstract

The tumor suppressor BRCA1-associated protein 1 (BAP1) is frequently mutated in uveal melanoma, where its loss is associated with poor prognosis. Although BAP1 has been implicated in homologous recombination (HR), its role in non-homologous end-joining (NHEJ) remains poorly defined. Here, we show that BAP1 functions as a central regulator of DNA double-strand break (DSB) repair by coordinating HR and NHEJ. BAP1 depletion disrupts recruitment and activity of the NHEJ machinery. Mechanistically, this defect is driven by aberrant accumulation of H2AK119-ub at DSB sites, promoting excessive DNA end resection and suppressing NHEJ activation. Importantly, inhibition of DNA end resection or suppression of H2AK119-ub restores NHEJ factor recruitment, establishing a causal link between BAP1-regulated histone modifications and repair pathway choice. Clinically, BAP1 loss correlates with genomic instability, providing a mechanistic basis for its association with poor outcomes in uveal melanoma. Collectively, these findings identify BAP1 as a gatekeeper of DSB repair fidelity, revealing a previously unrecognized role in safeguarding NHEJ and maintaining balanced DNA repair.

## Introduction

The ability of cells to sense and repair DNA double-strand breaks (DSBs), induced by endogenous or exogenous stressors such as chemotherapeutic agents or environmental insults, is essential for maintaining genome integrity(Ceccaldi *et al*, 2016). These highly toxic lesions interfere with vital processes including DNA replication and transcription, and, if left unrepaired or misrepaired, can lead to genomic alterations that profoundly affect cellular behavior (Jackson & Bartek, 2009). The accurate execution and proper choice between the two main DSB repair pathways, non-homologous end-joining (NHEJ) and homologous recombination (HR), are tightly regulated by histone post-translational modifications(Ceccaldi & Cejka, 2025). These modifications dynamically remodel chromatin to recruit the appropriate repair factors to DNA damage sites (Cejka & Symington, 2021; Clouaire & Legube, 2019). For example, methylation of Histone H4 at lysine 20 (H4K20me) and ubiquitylation of Histone H2A at lysine 15 (H2AK15ub) at DSB-flanking nucleosomes promote 53BP1 recruitment and downstream effectors that suppress DNA-end resection, thereby facilitating NHEJ (Cejka & Symington, 2021; Fradet-Turcotte *et al*, 2013). Conversely, ubiquitylation of Histone H2A at lysine 127 (H2AK127ub) by the BRCA1/BARD1 E3 ligase complex promotes HR (Densham *et al*, 2016). Intriguingly, monoubiquitylation of Histone H2A at lysine 119 (H2AK119ub) has also emerged as a regulator of repair pathway choice: Polycomb Repressive Complex 1 (PRC1)-mediated H2AK119ub deposition at DSBs support recruitment of the resection factor CtIP, thereby initiating end resection and promoting HR (Fitieh *et al*, 2022).

The tumor suppressor BRCA1-associated protein 1 (BAP1) is a nuclear ubiquitin carboxy-terminal hydrolase (UCH) that functions as the catalytic subunit of the Polycomb repressive deubiquitinase (PR-DUB) complex (Conway *et al*, 2021; Masclef *et al*, 2021). By removing H2AK119ub deposited by PRC1, BAP1 counteracts PRC1 activity, thereby reshaping chromatin landscapes, transcriptional programs, and key cellular processes including cell cycle progression, metabolism and cell death (Carbone *et al*, 2020). BAP1 has been implicated in the DNA damage response: it is phosphorylated by ATM upon DNA damage and localizes to break sites. BAP1-deficient cells display impaired HR and defective DSB repair (Ismail *et al*, 2014; Yu *et al*, 2014). However, the potential role of BAP1 in regulating NHEJ activation and execution has remained unexplored.

BAP1-associated tumorigenesis shows a distinctive spectrum, with a high frequency of mutations in relatively rare cancers such as uveal melanoma and malignant mesothelioma (Carbone *et al*., 2020; He *et al*, 2019). Uveal melanoma is the most common primary intraocular malignancy in adults, characterized by high lethality due to its propensity to metastasize, most often to the liver. Approximately 50% of patients develop metastases, with a median survival of ∼12 months following diagnosis (Krantz *et al*, 2017). While radiotherapy and enucleation can control primary tumors, no effective systemic therapies exist for metastatic uveal melanoma (Chua *et al*, 2021). BAP1 mutations are frequent in uveal melanoma, and its inactivation strongly correlates with metastatic dissemination and poor prognosis(Decatur *et al*, 2016; Karlsson *et al*, 2020). Understanding the molecular mechanisms by which BAP1 loss promotes tumor progression and therapy resistance in UM is therefore of urgent clinical importance.

In this study, we demonstrate that BAP1 loss associates with genomic instability in uveal melanoma. Mechanistically, we show that BAP1 deficiency impairs not only HR but also NHEJ, rendering cells unable to efficiently repair DSBs. In particular, BAP1 depletion induces aberrant DNA end resection in G1-phase cells, thereby antagonizing NHEJ. These findings establish BAP1 as a central regulator of DSB repair pathway choice and, importantly, reveal for the first time a mechanistic link between BAP1 activity and NHEJ regulation, with direct implications for genomic stability and therapeutic vulnerabilities in uveal melanoma.

## Results

### BAP1 mutations predict poor prognosis in uveal melanoma

The BAP1 gene locus is predominantly altered in specific tumors associated with the BAP1 tumor predisposition syndrome, including uveal melanoma, mesothelioma, intrahepatic cholangiocarcinoma (ICC), and kidney renal clear cell carcinoma (ccRCC), with alteration frequencies spanning from 10% to more than 35%, compared with only ∼3% across all human cancers (**Figure 1A**). In uveal melanoma, patients carrying BAP1 mutations exhibit significantly reduced overall survival (**Figure 1B, Supplemental figure S1A, S1B**), shorter disease-free survival (**Figure 1C**), and a greater prevalence of additional adverse prognostic markers (**Figure 1D**). Interestingly, this prognostic effect was not observed in individuals with malignant mesothelioma (**Supplemental Figure S1C**).

**Figure 1.**
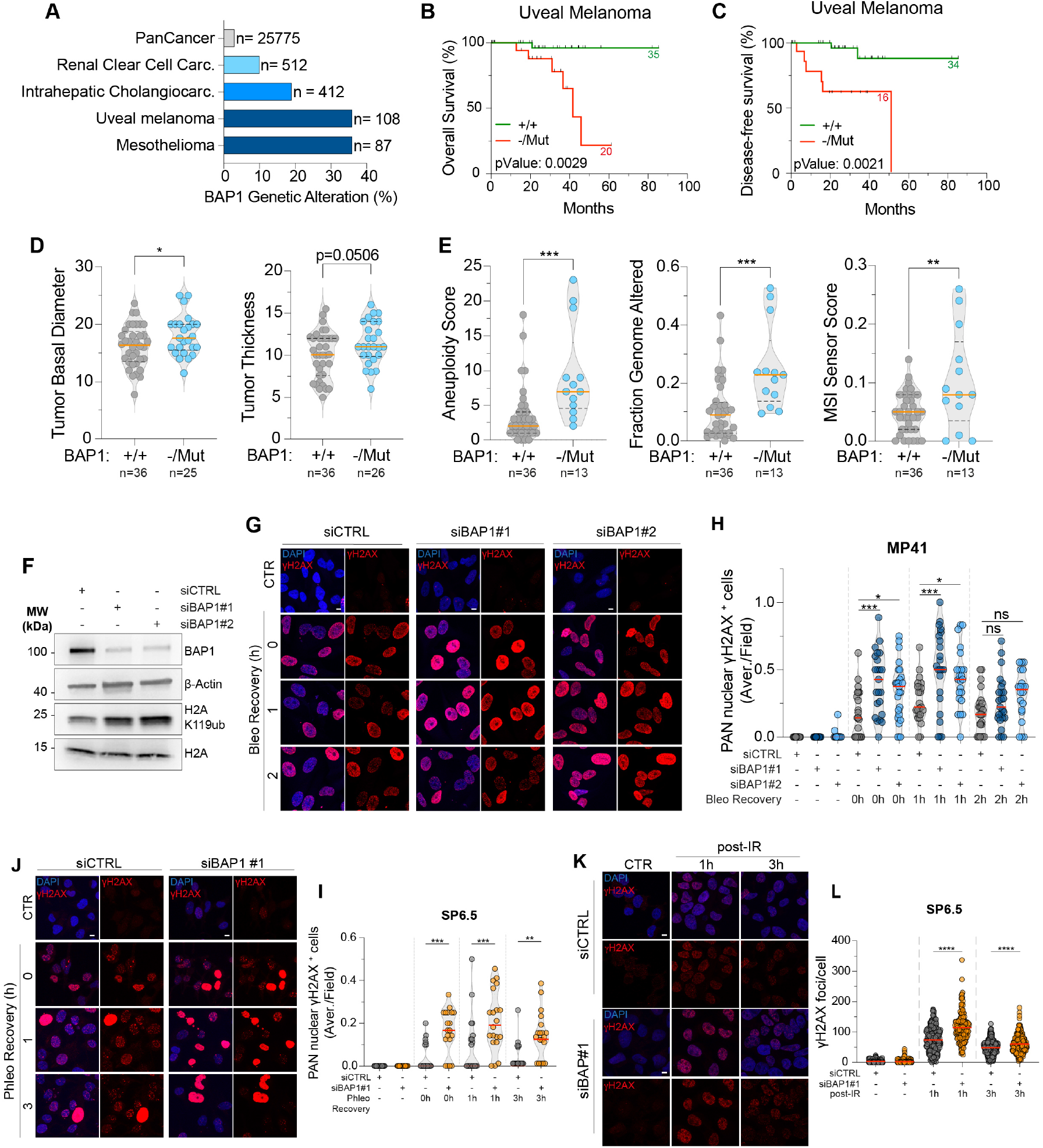
BAP1 mutations in uveal melanoma (UM) are associated with genomic instability. (A) Frequency of BAP1 genetic alterations in tumors from BAP1 cancer predisposition syndrome and in pan-cancer cohorts of metastatic tumors. Patient cohorts were retrieved from QIMR Oncotarget 2016(Johansson *et al*., 2016) and TCGA Firehose Legacy (UM), TCGA PanCancer Atlas (mesothelioma and kidney renal clear cell carcinoma)(Ellrott *et al*., 2018; Hoadley *et al*., 2018; Taylor *et al*., 2018), MSK Hepatology 2021 (intrahepatic cholangiocarcinoma)(Boerner *et al*., 2021), and MSK Cell 2021 (pan-cancer)(Nguyen *et al*., 2022) datasets via the cBioPortal platform. Data are shown as bar plots. (B,C) Kaplan–Meier survival curves for overall survival (B) and disease-free survival (C) in UM patients with diploid BAP1 wild-type (+/+, green) or BAP1 null/mutated (red) tumors. P-values are indicated. Data were retrieved from TCGA Firehose Legacy. (D, E) Violin plots showing tumor basal diameter, tumor thickness (D), aneuploidy score, fraction of genome altered and microsatellite instability (MSI) score (E) in diploid BAP1 wild-type versus BAP1 null/mutated UM patients. Orange line: median. *\*\*\*p<0*.*001 **p < 0*.*01, *p < 0*.*05*. P-values were calculated using an unpaired t-test. Data were retrieved from TCGA Firehose Legacy (D) and TCGA PanCancer (E). (F) Western blot analysis of BAP1 and H2AK119ub levels in BAP1-depleted MP41 cells. (G, I, K) Representative immunofluorescence images of γH2AX (red) at the indicated recovery time points after drug removal in MP41 (G) and SP6.5 (I) or post IR (5 Gy) (K). DAPI stains DNA (blue). Scale bar: 10 µm. (H,J) Violin plots quantifying the percentage of pan-nuclear γH2AX+ cells per field across 3 independent experiments in MP41 (H) and 2 independent experiments in SP6.5 (J). Each dot represents a field. Red line: median. *\*\*\*p < 0*.*001, **p < 0*.*01, *p < 0*.*05, ns: not significant*. P-values were determined by Ordinary one-way ANOVA. (L) Scatter dot plot showing the quantification of γH2AX foci in SP6.5 following IR. 2 independent experiments were conducted. Red line: Mean. *\*\*\*\*p < 0*.*0001*. P-values were determined by Ordinary one-way ANOVA.

BAP1 has been broadly implicated in maintaining genome integrity, in part through its role in HR–mediated repair of DSBs. We therefore investigated whether BAP1 mutational status correlates with genomic instability. In uveal melanoma, BAP1 mutated tumors displayed significantly higher levels of aneuploidy, fraction of the genome altered and microsatellite instability (MSI) scores, compared with wild-type diploid tumors (**Figure 1E**). This association, however, was not evident in other tumor types characterized by frequent BAP1 alterations (**Supplemental Figure S1D-S1G**). Together, these findings suggest that BAP1 mutations drive distinct mechanisms of genomic instability in uveal melanoma, which may underlie the prognostic impact of BAP1 mutations.

### BAP1 loss compromises DSB repair in uveal melanoma cells

We next investigated the molecular mechanisms underlying the compromised genome integrity observed in uveal melanocytes upon loss of BAP1. For this purpose, we used the BAP1-proficient human uveal melanoma cell lines MP41 and SP6.5. Because stable CRISPR/Cas9-mediated deletion of BAP1 was incompatible with *in vitro* survival of these cells, we reduced BAP1 expression using short interfering RNAs (siRNAs). As expected, BAP1 depletion led to the accumulation of the H2AK119ub histone mark, a well-established post-translational modification dynamically regulated by BAP1 deubiquitinase activity (**Figure 1F**) (Szczepanski & Wang, 2021).

We next evaluated the impact of BAP1 loss on the resolution kinetics of DSBs following genotoxic stress. BAP1-deficient cells displayed impaired clearance of radiomimetic-induced DNA damage over time: immunofluorescence and Western Blot analyses for γH2AX showed an increased proportion of γH2AX signal during the recovery phase in BAP1-silenced MP41 (**Figure 1G,H, Supplemental figure S2A, S2C**) and SP6.5 (**Figure 1I-J, Supplemental figure S2B**) cells compared with controls. Consistent with these findings, ionizing radiation (IR) induced a more pronounced yH2AX foci formation in BAP1-depleted SP6.5 uveal melanocytes (**Figure 1K, 1L, Supplemental Figure S2D**). Together, these results demonstrate that BAP1 loss impairs DSB repair capacity, thereby compromising the maintenance of genome stability.

To further dissect the mechanistic role of BAP1 in double-strand break (DSB) repair in uveal melanoma, we examined the recruitment of key upstream repair factors from both the homologous recombination (HR) and non-homologous end joining (NHEJ) pathways following DSB induction. As expected(Ismail *et al*., 2014; Yu *et al*., 2014), BAP1-depleted cells exhibited a pronounced defect in RAD51 foci formation compared with BAP1-proficient cells, immediately after bleomycin removal and recovery (0 and 1h) (**Figure 2A-C**). Consistent with this, RAD51 was efficiently recruited to damage sites at 3 h post-ionizing radiation (IR) in control MP41 cells, but failed to accumulate in both BAP1- and BRCA1-depleted cells **(Figure 2D-F, Supplemental Figure S3A**).

**Figure 2.**
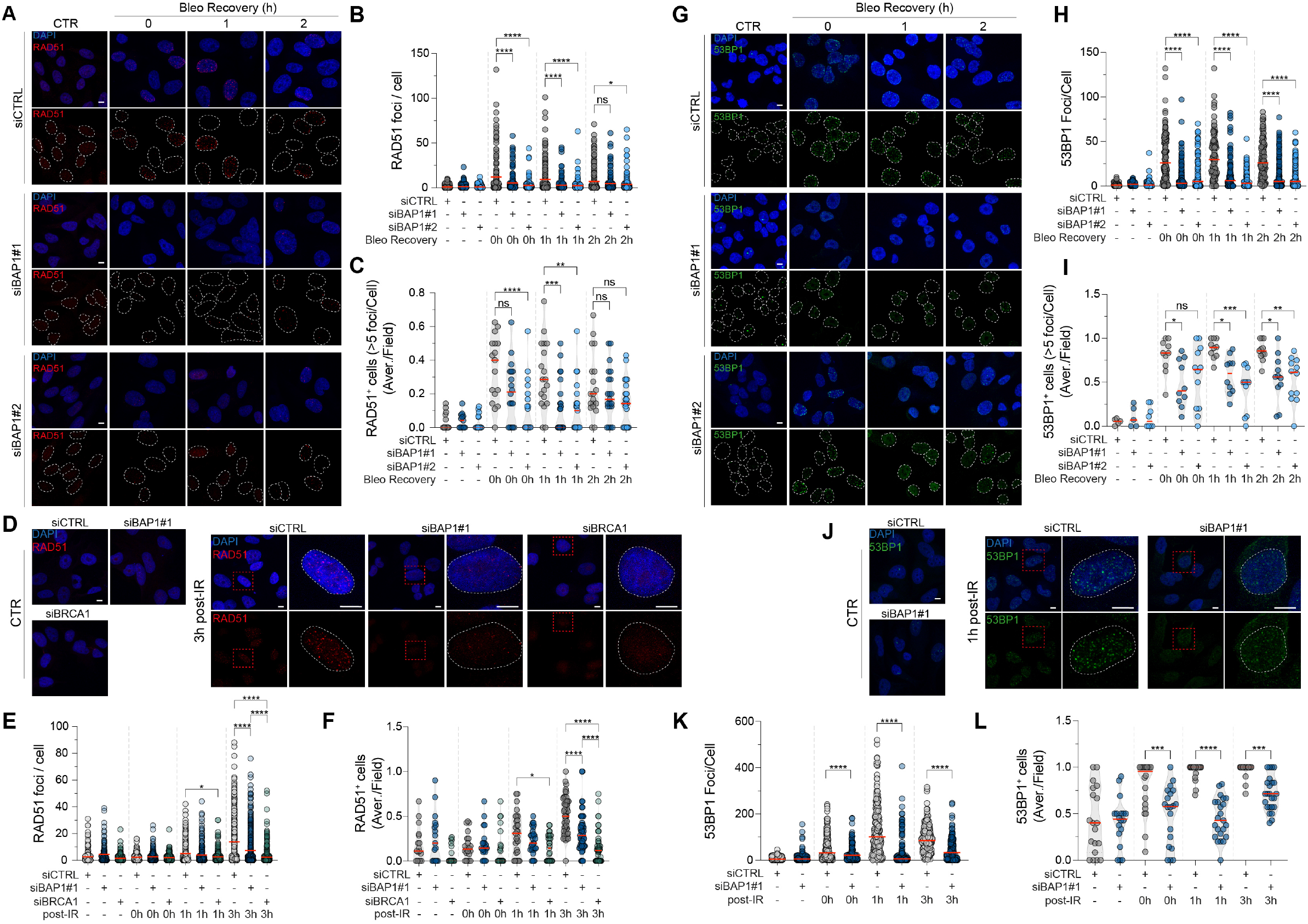
BAP1 depletion impairs the recruitment of HR and NHEJ repair factors. (A,D) Representative confocal images of RAD51 (red) in MP41 cells transfected with siCTRL or siBAP1 at the indicated recovery time points from bleomycin treatment (A) or following 5 Gy IR (D). DAPI stains DNA (blue). Scale bar: 10 µm. (B, C, E, F) Scattered plot showing the number of RAD51 foci per cell (B, E) and violin plot showing the proportion of RAD51+ cells (>5 foci/nucleus) (C,F) over the total cells per field (represented as dots) in 3 (B,C) and 2 (E,F) independent experiments. Red line: mean (B, E), median (C, F). (G, J) Representative confocal images of 53BP1 (green) in MP41 cells transfected with siCTRL or siBAP1 at the indicated recovery time points from bleomycin treatment (G) or following IR (J). DAPI stains DNA (blue). Scale bar: 10 µm. (H,I,K,L) Scattered plots showing the number of 53BP1 foci per cell (H,K) or violin plots showing the proportion of RAD51+ cells (>5 foci/nucleus) (I,L) over the total cells per field (shown as dots) in 3 (H,I) and 2 (K,L) independent experiments. *\*\*\*\*p < 0.0001, ***p < 0.001, **p < 0*.*01, *p < 0*.*05, ns: not significant*. P-values were determined by Ordinary one-way ANOVA.

We next analyzed the recruitment of 53BP1, a critical regulator of DSB repair pathway choice that inhibits DNA end-resection and promotes NHEJ(Cejka & Symington, 2021). Unexpectedly, BAP1-depleted MP41 and SP6.5 cells displayed also a drastic reduction in 53BP1 foci formation during recovery following 24 h radiomimetic treatment (**Figure 2G-I, Supplemental Figure S3B-S3D**). Similar trend was also observed in the response of BAP1-deficient cells to IR treatment, where they failed to engage 53BP1 loading at the damage post IR (**Figure 2J-L, Supplemental Figures S3E-S3G)**.

Together, these results indicate that BAP1 loss impairs the recruitment of both RAD51 and 53BP1 to DSBs, thereby disrupting key steps of both HR and NHEJ repair pathways.

### BAP1 is required for execution of both HR and NHEJ DNA repair pathways

Based on the observation that markers of both HR and NHEJ were altered upon BAP1 silencing, we next sought to verify the functional efficiency of these DSB repair pathways upon BAP1 loss.

To this end, we employed stably integrated reporter plasmids that allow quantification of HR and NHEJ activity. Specifically, we used the HR-EGFP/5′EGFP (**Figure 3A**) and HR-EGFP/3′EGFP (**Supplemental Figure S5A**) reporters to monitor HR (Akyuz *et al*, 2002), along with the PIMEJ5GFP reporter system to assess NHEJ (**Figure 3D**)(Bennardo *et al*, 2008). Notably, the HR-EGFP/5′EGFP reporter detects exclusively conservative HR, whereas the HR-EGFP/3′EGFP reporter captures both conservative HR and single-strand annealing (SSA). Both HR reporters contain a mutated EGFP gene harboring an I-SceI restriction site, along with either a 3′EGFP or 5′EGFP donor sequence in tandem, which serves as a template for repair by HR. In contrast, the PIMEJ5GFP reporter contains two distal tandem I-SceI recognition sites flanking both a puromycin resistance cassette and the EGFP gene. Expression of I-SceI and its subsequent nuclear translocation induces site-specific DSBs within the reporters. Restoration of EGFP fluorescence occurs only when the break is successfully repaired, via HR in the HR reporters (HR-EGFP/5′EGFP or HR-EGFP/3′EGFP), or via NHEJ in the PIMEJ5GFP system.

**Figure 3.**
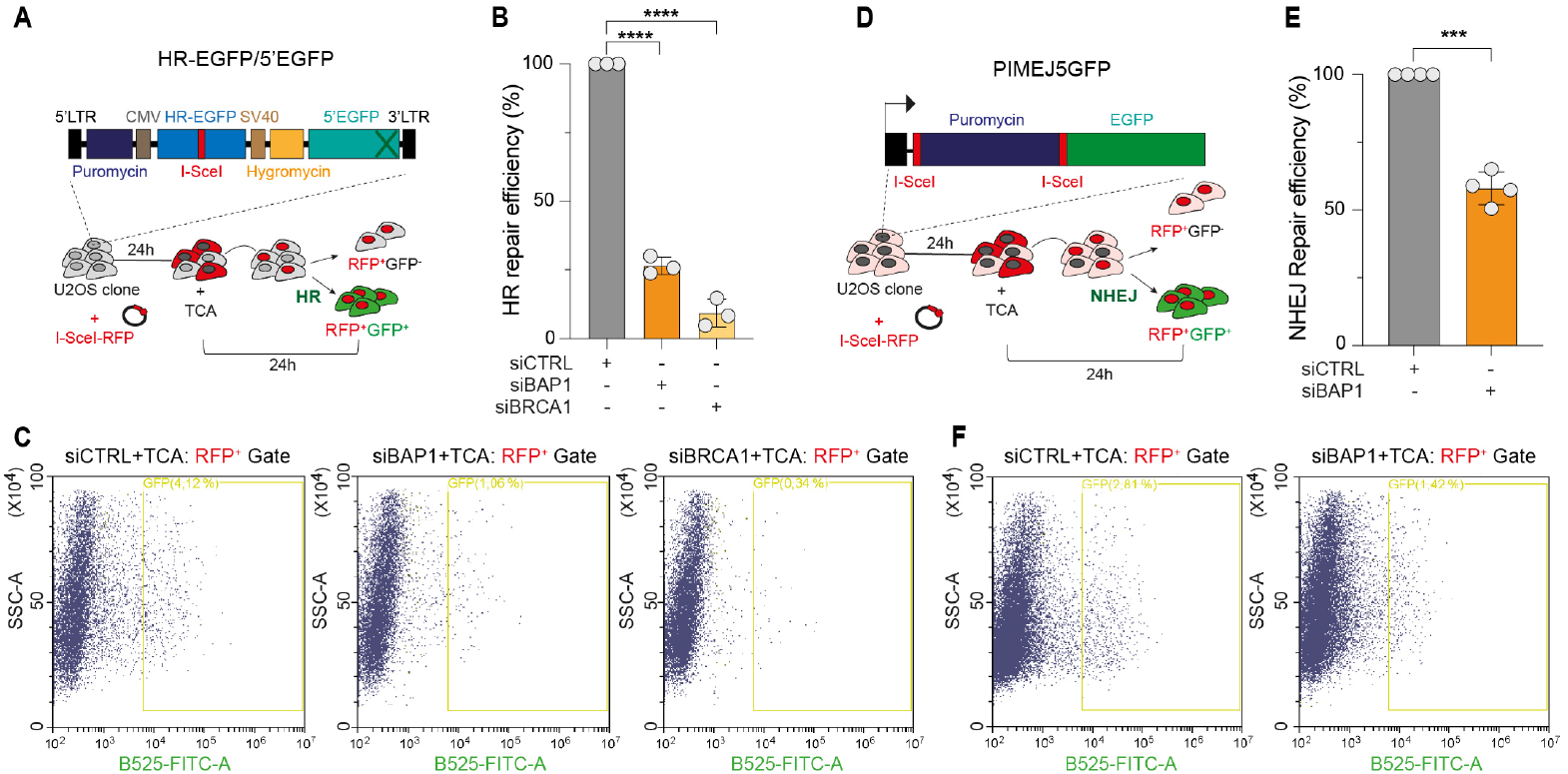
Loss of BAP1 compromises HR and NHEJ repair efficiency. (A, D) Schematics of the HR-EGFP/5′EGFP (A) and PIMEJ5GFP (D) reporter constructs. HR-EGFP/5′EGFP includes a mutated EGFP gene containing an I-SceI recognition site and a truncated 5′EGFP sequence serving as a repair template. PIMEJ5GFP contains two distal I-SceI sites flanking a puromycin gene and the EGFP sequence. Stable U2OS clones with chromosomally integrated reporters were transfected with siCTRL, siBAP1, or siBRCA1. The following day, cells were transfected with I-SceI-RFP (red fluorescence) and treated with triamcinolone acetonide (TCA) 24 h later to induce nuclear translocation of I-SceI. RFP+ cells were analyzed for GFP fluorescence. (B,E) Bar plots quantifying HR (B) and NHEJ (E) repair efficiency across ≥3 independent experiments. Data normalized to control and presented as mean ± SD. ****p < 0.0001. P-values by Ordinary one-way ANOVA (B) or paired t-test (E). (C, F) Representative flow cytometry plots showing GFP+ cells (yellow gate) within the RFP+ population under each condition. GFP fluorescence: B525-FITC-A; side scatter: SSC.

Analysis of HR-EGFP/5′EGFP cells showed that repair capacity (RFP^+^GFP^+^ population) was markedly reduced to 26.5% upon BAP1 depletion (**Figure 3B,C; Supplemental Figures S4A,S4B**). A similar reduction was observed in the HR-EGFP/3′EGFP clone (**Supplemental Figures S5B-S5D**). Remarkably however, we also found that NHEJ repair efficiency decreased to 58% in siBAP1-depleted cells (**Figures 3E, 3F; Supplemental Figures S4C, S4D**), consistent with the defective 53BP1 localization of BAP1 deficient cells. Together, these results indicate that BAP1 loss impairs not only HR but also the NHEJ double-strand break repair pathway.

### BAP1 loss triggers aberrant DNA end resection

The ubiquitination of histone H2A at lysine 119 (H2AK119ub) at DSB sites influences the activation of DNA end resection and consequently the execution of HR(Fitieh *et al*., 2022). As BAP1 is responsible for erasing H2A-K119ub, we hypothesized that BAP1 loss might underlie aberrant DNA end resection in uveal melanoma cells following DSB, thus impacting the ability to initiate NHEJ also in G1 phase. BAP1 depletion caused an accumulation of H2AK119ub in G1 cells untreated and exposed to X-ray (**Figure 4A, Supplemental Figure S6A**). This regulation appeared specific as other histone modifications involved in DSB repair execution were not altered (**Supplemental Figure S6B)(Ramachandran *et al*, 2016)**. To test the direct involvement of H2AK119ub at DSB sites, we carried out chromatin immunoprecipitation analysis (ChIP-qPCR) using the DIvA inducible cell system. Following AsiSi-induced DSBs, we detected a substantial enrichment of H2AK119ub at three canonical AsiSI sites (Chr22, Chr6 and Chr1)(Iacovoni *et al*, 2010) (**Figure 4B,C, Supplemental figure S6C**). Notably, DNA damage-dependent elevation of H2AK119ub was further increased by BAP1 depletion, consistent with a direct role at DSB sites and a regulation mediated by BAP1 (**Figure 4C**).

**Figure 4.**
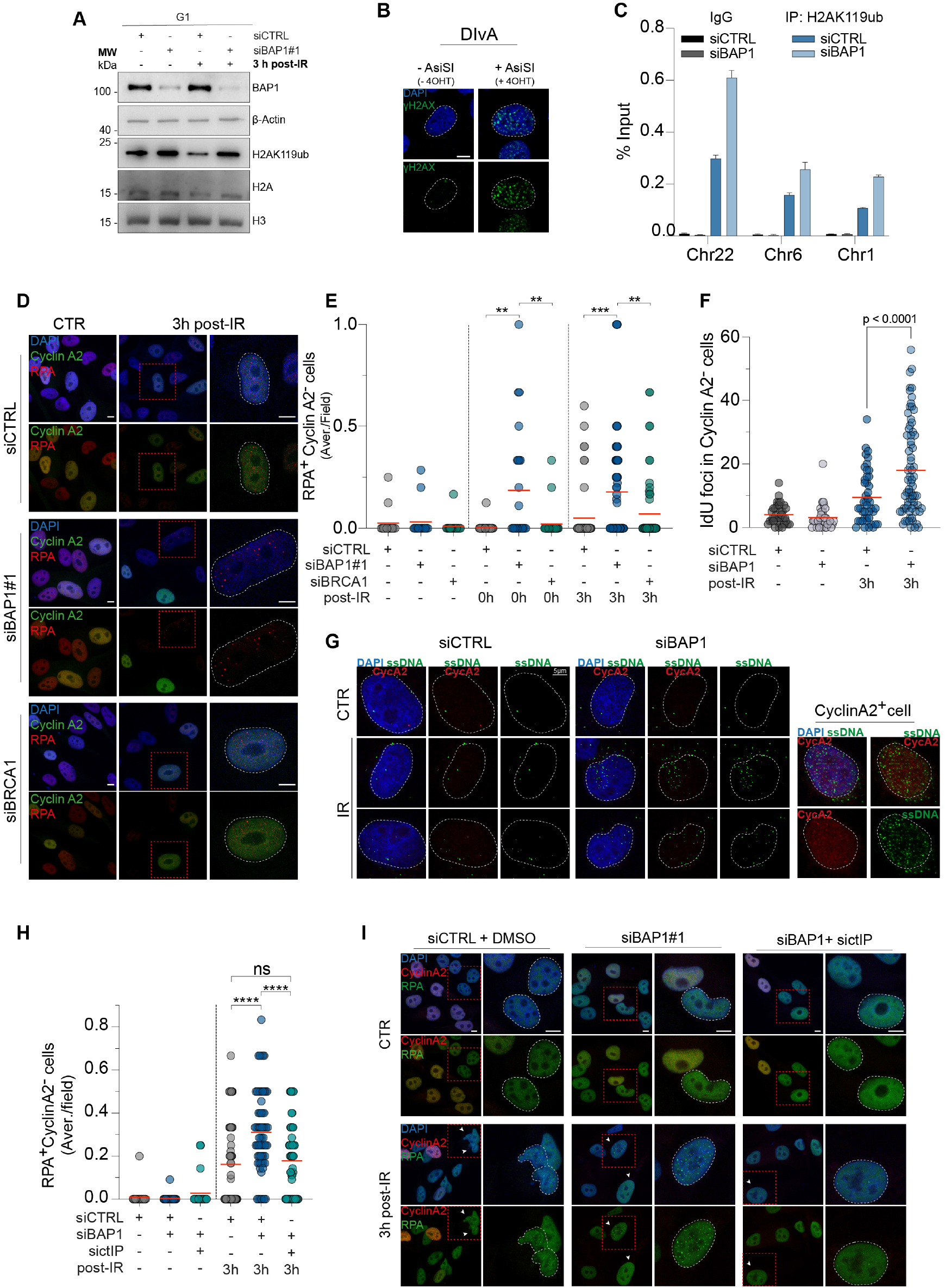
BAP1 depletion promotes aberrant DNA end-resection in G1 phase. (A) Western blot analysis of H2AK119ub levels upon BAP1 depletion in G1 cells. MP41 were irradiated (5 Gy) at 6 hours from Nocodazole release and collected 3 hours post IR. (B) Representative immunofluorescence images showing AsiSI-induced DSB via γH2AX foci detection (green) after 4 hours following the addition of 300 nM 4-OHT. DAPI stains nuclei. Scale bar 10 µm. C) Representative ChIP-qPCR showing the accumulation of H2AK119ub at the AsiSI-induced DSB sites following 4-OHT treatment (4h) in BAP1-depleted DIvA cells (data shown as mean + SD of technical triplicates). Two independent experiments were conducted. (D,E) Representative immunofluorescence images for cyclin A2 (green), RPA (red) co-staining in MP41 cells post IR following BAP1 depletion (D). Scatter dot plot showing the number of RPA^+^ Cyclin A2^−^ cells (G1 phase) over the total fraction of CyclinA2-cells indicated as average/field (E). Red line: Mean. **p < 0.01, P-values by Ordinary one-way ANOVA. (F,G) Non-denaturing IdU labeling assay. Quantification of IdU foci in Cyclin A2^-^ cells following IR (F). Two independent experiments were conducted. Red line: Mean. P-values by Ordinary one-way ANOVA. Representative immunofluorescence images are shown in panel G. DAPI stains nuclei, Scale bar 10 µm. (H,I) Scatter dot plot quantifying the number of RPA^+^ Cyclin A2^−^ cells (G1 phase) in MP41 cells co-depleted of BAP1 and ctIP. Red line: Mean. ****p < 0.0001, p-value by 2way Anova. Representative immunofluorescence images are shown in I. Scale bar 10 µm.

To test the hypothesis that altered regulation of H2AK119ub at DSBs upon BAP1 loss promotes aberrant DNA end resection, we analyzed RPA recruitment as a marker of end resection. Consistent with this hypothesis, BAP1 depletion increased RPA foci formation during DSB recovery (**Supplemental Figure S6D**), indicating dysregulated control of DNA end resection. We next examined RPA foci formation in Cyclin A2–negative (G1) cells to determine whether BAP1 loss induces unscheduled end resection in G1, potentially explaining the NHEJ defect. This analysis revealed that BAP1 loss led to a substantial accumulation of RPA foci (**Figure 4D, E and Supplemental Figure S6E, S6F**). To further confirm the occurrence of unscheduled DNA end resection, we performed a non-denaturing IdU labeling assay, which revealed increased single-stranded DNA (ssDNA) generation upon DSB induction in Cyclin A2–negative cells following BAP1 depletion (**Figure 4F, G and Supplemental Figure S7A**). Together, these results indicate that BAP1 loss promotes unscheduled DNA end resection in G1 cells.

To establish a causal link between aberrant end resection and defective NHEJ, we abrogated end resection by silencing its key regulator, CTBP-interacting protein CtIP(Nicolas *et al*, 2024; Zhao & Zhang, 2025). Immunofluorescence analysis in Cyclin A2–negative cells showed that CtIP silencing reverted the aberrant RPA accumulation observed in BAP1-depleted cells, restoring RPA foci levels to those seen in control cells (**Figure 4H, I and Supplemental Figure S7B**). Together, these data demonstrate that BAP1 loss triggers unscheduled DNA end resection and RPA recruitment in G1 cells, which likely prevents NHEJ activation and impairs resolution of DSBs. Together, these data demonstrate that BAP1 loss triggers unscheduled DNA end resection and RPA recruitment in G1 cells, potentially explaining the altered NHEJ activation.

### H2AK119ub accumulation in BAP1-depleted cells counteracts NHEJ by promoting DNA end resection

To establish the mechanistic basis of altered NHEJ activation in BAP1-deficient cells, we sought to define a causal link between H2AK119ub accumulation, aberrant DNA end resection, and NHEJ execution. To this end, we employed a pharmacological approach using PTC-209, a small-molecule inhibitor that reduces H2AK119 ubiquitylation by targeting BMI1, a catalytic component of the PRC1 complex(Kreso *et al*, 2014). We first verified that PTC-209 efficiently abolished the increase in H2AK119ub at DSB sites induced by BAP1 depletion (**Figure 5A and Supplemental Figure S7C, D**). Next, cell cycle–resolved analysis of RPA recruitment revealed that inhibition of H2AK119ub accumulation by PTC-209 reduced RPA loading in G1-phase (Cyclin A2–negative) cells (**Figure 5B–D and Supplemental Figure S7E**), indicating that excessive H2AK119ub accumulation drives aberrant DNA end resection. To determine whether this dysregulation contributes to defective NHEJ activation in BAP1-depleted G1 cells, we measured 53BP1 recruitment following IR in cells proficient or deficient for BAP1 in the presence of PTC-209. As predicted, PTC-209 restored 53BP1 loading in BAP1-deficient cells (**Figure 5E, 5F**). Together, these data establish a causal relationship between H2AK119ub accumulation in BAP1-deficient G1 cells, aberrant end resection (RPA recruitment), and impaired NHEJ.

**Figure 5.**
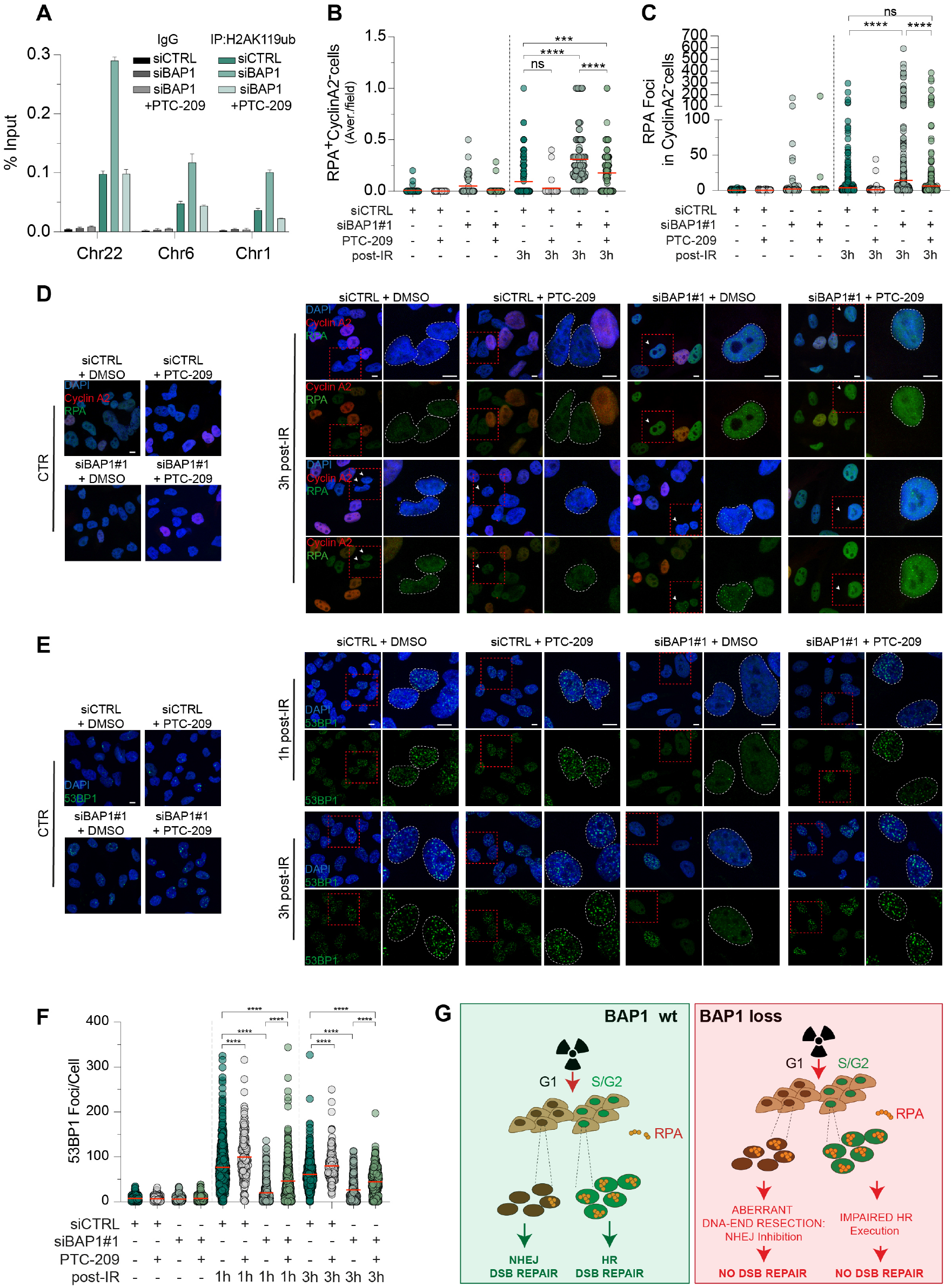
PTC-209 partially rescues NHEJ defects in BAP1-depleted cells. (A) Representative ChIP-qPCR showing the enrichment of H2AK119ub at the AsiSI-induced DSB sites following PTC-209 (10 µM 18h) and 4-OHT (4H) treatments in BAP1-depleted DIvA cells (data shown as mean + SD of technical triplicates). Two independent experiments were conducted. (B-D) Scatter dot plots showing the number of RPA^+^ Cyclin A2^−^ cells over the total fraction of CyclinA2^-^ cells per field (B) and RPA foci in Cyclin A2^−^ cells (C) at 3 h post-IR in BAP1-depleted MP41 cells ± PTC-209 treatment. Red line: mean. ****p < 0.0001, ***p < 0.001, ns: not significant. P-values by Ordinary one-way Anova. Representative immunofluorescence images of RPA (green) and Cyclin A2 (red) (D). DAPI stains DNA. MP41 were treated with PTC-209 (0.1 µM, 48 h) prior to irradiation and collected at 3 h post-IR. Arrows mark Cyclin A2^−^ (G1 phase) cells with RPA foci. Scale bar: 10 µm. (E,F) Representative immunofluorescence images of 53BP1 (green) in BAP1-depleted MP41 cells ± PTC-209 treatment at 1 and 3 h post-IR. DAPI stains DNA. Scale bar: 10 µm (E). Scatter dot plot quantifying 53BP1 foci per cell. Red line: mean. ****p < 0.0001. P-values by 2way ANOVA (F). (G) Schematic model. In BAP1-proficient cells, BAP1 deubiquitylates H2AK119ub deposited by PRC1, permitting 53BP1 recruitment and NHEJ execution in G1. In BAP1-depleted cells, H2AK119ub accumulates at the DSB sites, blocking 53BP1 loading, and driving aberrant DNA end resection with RPA recruitment in G1, where HR is not available.

Collectively, these findings highlight that BAP1-mediated regulation of histone ubiquitylation at DSBs is essential for proper engagement of NHEJ in G1 cells and for maintaining appropriate DSB repair pathway choice.

## Discussion

Our study uncovers a previously unrecognized role for BAP1 in regulating NHEJ DSB repair, with direct implications for the genomic instability and aggressive clinical behaviour of uveal melanoma.

While BAP1 has been linked to HR through earlier reports (Ismail *et al*., 2014; Yu *et al*., 2014), our findings establish that its function extends beyond HR, encompassing a crucial role in NHEJ. Specifically, we show that BAP1 loss triggers aberrant DNA end resection in G1 cells, thereby antagonizing NHEJ and leaving cells unable to effectively resolve DSBs. This dual impairment of both HR and NHEJ provides a mechanistic explanation for the pronounced genomic instability associated with BAP1 mutations in uveal melanoma (**Figure 5G**).

Histone post-translational modifications are well-established regulators of DSB repair pathway choice, acting as dynamic chromatin signals that guide the recruitment of repair factors (Clouaire & Legube, 2019). Our work highlights the importance of H2AK119 ubiquitylation as a determinant of pathway engagement. We demonstrate that, in the absence of BAP1, accumulation of H2AK119ub at DSB sites drives unscheduled recruitment of the resection factor RPA in G1-phase cells that can be reverted by depleting ctIP. This inappropriate activation of end resection prevents 53BP1 loading, thus impairing NHEJ, a repair pathway canonically required during G1 when homologous templates are unavailable. Importantly, pharmacological inhibition of H2AK119ub deposition at the DSB with PTC-209 restored 53BP1 accumulation and reduced aberrant resection, establishing a direct causal link between BAP1-regulated deubiquitylation, chromatin remodelling at DSBs, and pathway choice.

These findings position BAP1 alongside other chromatin regulators that control the fidelity of DSB repair pathway choice. BAP1 preserves NHEJ by ensuring proper initiation of DNA-end resection, while also supporting HR by enabling its successful completion. Cells lacking BAP1 exhibit concurrent defects in both pathways: NHEJ is compromised due to aberrant end resection, and HR fails to proceed, as evidenced by the absence of RAD51 loading. Thus, BAP1 acts as a central “gatekeeper” of pathway choice and execution, ensuring that DSBs are repaired by the appropriate mechanism in a context-dependent manner. The simultaneous impairment of HR and NHEJ in BAP1-deficient cells creates a distinct state of “repair exhaustion,” which may promote the accumulation of structural alterations and aneuploidy observed in BAP1-mutant uveal melanoma tumors.

From a clinical perspective, these findings provide a mechanistic explanation for the aggressive phenotype and poor prognosis associated with BAP1-mutant uveal melanoma. Moreover, our analysis of cancer patients indicates that BAP1 inactivation in uveal melanoma is driven exclusively by single-nucleotide alterations, including substitutions and nonsense mutations, which ultimately result in loss-of-function effects(Bhattacharya *et al*, 2015; Bott *et al*, 2011). In contrast, other BAP1-inactivated tumors exhibit a more balanced distribution of mutations and deletions. Our findings also suggest potential therapeutic vulnerabilities. On one hand, defective HR renders these tumors potentially sensitive to PARP inhibitors or other agents that exploit HR deficiency. On the other hand, the unexpected impairment of NHEJ may influence responses to radiotherapy or genotoxic chemotherapy, as cells lacking both major repair pathways might accumulate lethal damage but could also activate compensatory error-prone mechanisms. Targeting H2AK119ub dynamics, as demonstrated with BMI1 inhibition, may represent a novel therapeutic strategy to restore repair pathway balance or exploit vulnerabilities specific to BAP1-deficient cancers.

In summary, our work identifies BAP1 as a critical regulator of DSB repair pathway choice. By restraining H2AK119ub-driven end resection, BAP1 ensures proper engagement of NHEJ, thereby maintaining genomic stability. Loss of this regulation simultaneously disables HR and NHEJ, fuelling genome instability and tumour progression in uveal melanoma. These findings not only advance the understanding of BAP1 biology but also open new avenues for therapeutic intervention in BAP1-deficient tumours.

## Acknowledgments

The authors would like to thank Dr. Halime Kalkavan for kindly providing the SP6.5 uveal melanoma cell line(Soulieres *et al*, 1991) and Dr. Gaelle Legube for kindly providing the DiVa cell model(Iacovoni *et al*., 2010). This work has been supported by the DFG to IA, (under the TRR353 “Death Decision” projects A05), the Carl Zeiss Stiftung to IA (Endowed Professorship, #15972218, 2022-2027; Prisma Programme, #P2022-5-003), by the cooperation between Carl Zeiss Stiftung and German Scholars Organization with the Fund for international researchers to IA (#15978021) and by the MWK funding to IA. (Az. MWK31-7532-522/1/6).

## Data Availability

All the data are made available upon request.

## Materials availability

All unique/stable reagents generated in this study are available from the lead contact with a completed materials transfer agreement.

## Authors Contribution

Conceptualization, IA; methodology, SC, MRS; investigation, SC, CS, AB and NG; writing – original draft, SC; writing – review & editing, LW and IA; resources, LW and IA; supervision, LW and IA

## Material and Methods

### Cell culture and siRNA transfection

MP41 human cells were purchased from ATCC (MP41-CRL-3297 ™) and were maintained in RPMI medium (Gibco) supplemented with 20% fetal bovine serum (FBS), sodium pyruvate (1 mM), Hepes Buffer (10 mM) and penicillin–streptomycin (100 U/ml).

SP6.5 cells were kindly provided by Kalkavan Halime and were grown in RPMI medium (Gibco) supplemented with 10% fetal bovine serum (FBS) and penicillin–streptomycin (100 U/ml).

U2OS human cells were purchased from ATCC (HTB-96 ™ATCC) and were maintained in Dulbecco’s Modified Eagle Medium (DMEM) supplemented with 10% FBS and penicillin-streptomycin (100 U/ml).

DIvA cells were kindly provided by Gaelle Legube and were grown in Dulbecco’s Modified Eagle Medium (DMEM) supplemented with 10% FBS, penicillin-streptomycin (100 U/ml) and Puromycin (1 μg/ml). All cell lines were grown at 37°C, 5% CO2.

All cell lines were transfected with 20 nM of scramble (siCTRL) or BAP1, BRCA1, ctIP targeting siRNAs (see key sources table) using Lipofectamine RNAiMAX (Invitrogen, ThermoFisher Scientific) according to the manufacturer’s protocol. Collection was performed at 72 hours from siRNA transfection. For DNA damage induction, MP41 and SP6.5 transfected cells were treated with bleomycin (50 µg/ml) or phleomycin (50 µg/ml) for 24 hours or irradiated with 5 Gy or 10 Gy. For AsiSI induction, DIvA cells were treated with 300 nM (Z)-4-Hydroxytamoxifen (4-OHT Biozol, Cat# APE-B5421) for 4 hours before sample collection for ChIP-qPCR or immunofluorescence analysis.

For rescue experiments with BMI inhibition, BAP1-depleted MP41 cells were treated with DMSO or 0,1 µM PTC-209 (BMI-inhibitor, Sigma Aldrich) for 48 hours prior IR-induced DNA damage. Cells were then collected for immunofluorescence or western blot analysis at the indicated hours post drug removal or IR-induced DSBs. DIvA cells for ChIP-qPCR were treated with DMSO or 10 µM PTC-209 for18 hours prior AsiSI induction.

### Cell synchronization and Cell cycle Analysis

For cell synchronization, MP41 cells were incubated with 100 nM Nocodazole for 16 hours. Following mitotic shake-off, cells were allowed to recover in normal medium and irradiated (5 Gy) at 6 hours after nocodazole release. Cells were collected prior to irradiation for cell cycle analysis via Propidium iodide (PI) staining (100 μg/mL, O.N) and 3h post-IR for western blot analysis. Pi staining for both asynchronized and synchronized cells was acquired using the Beckman Coulter CytoFLEX™ and at least 10000 events were acquired per sample.

### Generation of stable cell lines with integrated reporters

For the generation of U2OS clones with stably chromosomally integrated DSB repair reporters, U2OS cells were transfected with 5 μg of the following plasmids, pimEJ5GFP, HR-EGFP/3’EGFP or HR-EGFP/5’EGFP using PEI reagent according to the manufacturers’ instructions.

U2OS transfected cells were maintained in complete RPMI medium supplemented with 2 μM of Puromycin. After Puromycin selection, isolation of individual cell clones for each repair plasmid was obtained by serial dilution. Single clones were then amplified and tested for EGFP fluorescence upon I-SCEI induction using flow cytometry (Beckman Coulter CytoFLEX™).

### DNA Damage Reporter Assays

HR-EGFP/3’EGFP, HR-EGFP/5’EGFP and Pim-EJ5-GFP U2OS clones were transfected with siRNA control or against BAP1 (Select BAP1#2 siRNA, s15822) or BRCA1 to assess repair efficiency respectively of homologous recombination (HR) and single-strand annealing (SSA), conservative HR (cHR) and total non-homologous end joining (NHEJ) DNA repair pathways.

All U2OS clones were transfected with 2 µg ISceI-GR-RFP plasmid and after 24 hours, treated with 0.4 µM triamcinolone acetonide (TCA) or DMSO. TCA-driven-I-SceI-RFP nuclear translocation led to a site-specific cleavage within the integrated EGFP DNA sequence. DNA repair efficiency for HR or NHEJ was evaluated by measuring the EGFP fluorescence signal upon the reconstitution of the functional EGFP coding sequence.

As a positive control for EGFP fluorescence, U2OS cells were co-transfected with wtEGFP and I-SceI-RFP plasmids as described previously. Sample were collected at 72 h post-silencing and 24 h post-TCA induction. EGFP and RFP fluorescence signals were detected by flow cytometric analysis using the Beckman Coulter CytoFLEX™ and at least 10000 events were acquired per condition. The percentage of Individual GFP-positive cells were then examined within the RFP-positive cell population.

### Protein Extraction and Western blot analysis

MP41 or U2OS cells were lysed using RIPA buffer supplemented with protease and phosphatase inhibitors (Sigma Aldrich). 20 µg of proteins were then denatured at 98°C for 10 minutes, resolved on a SDS– polyacrylamide gel and transferred onto an Amersham Hybond P0.45 PVDF membrane (Sigma Aldrich). Membranes were incubated with a blocking solution of 5% non-fat dried milk diluted in PBS 0.1% Tween-20 for 2h at RT. Incubation of primary antibodies was performed overnight at 4°C (see key resources table). Membranes were then incubated with the appropriate anti-mouse or anti-rabbit peroxidase-conjugated secondary antibodies (Bio-Rad) at room temperature for 1 h. Chemiluminescence signals were detected using ECL chemiluminescence kit (Thermofisher, cat# 32106X4) or SuperSignalTM West Dura Extended Duration Substrate (Thermofisher).

### RNA extraction, RT and qPCR

RNA extraction was performed using RNeasy Mini Kit (Qiagen) according to the manufacturer’s instruction. 1 µg of extracted RNA was successively reverse transcribed with the SensiFAST cDNA Synthesis kit (Biocat, cat# BIO-65054). Real time PCR (qPCR) was performed using PerfeCTa® SYBR® Green FastMix (VWR, Cat# 733-1390). BAP1 and TBP target genes were amplified using QuantStudio 1 RealTime PCR System (Applied Biosystems) (See key resources table). qPCR results were expressed as a relative gene expression and calculated following the 2−ΔΔct method after normalization to human TATA-binding protein (TBP).

### Chromatin immunoprecipitation (ChIP)

For ChIP experiments following AsISI-induced DNA damage, DiVA cells were fixed with 1% formaldehyde for 10 minutes at Room Temperature and the reaction was quenched with 0.125 M glycine for 5 minutes at RT. Then, extracted nuclei were sonicated for 10 cycles (30” ON - 30” OFF, High settings) at 4°C with a BioruptorTM UCD-200 Diagenode to obtain chromatin fragments of 250-600 bp. For each immunoprecipitation, 100 µg chromatin and 2 µg of antibody were used. The immunoprecipitation was carried out using Dynabeads Protein G (Invitrogen, cat# 10004D) at 4°C for 4 hours with gentle rotation. Then, the beads were washed twice with IP Dialysis buffer (50mM Tris-HCl pH 8.0, 2mM EDTA, 0.2% N-Laurylsarcosine) and 4 times with IP wash buffer (100mM Tris-HCl pH9.0, 500mM LiCl, 1% NP40, 1% NaDOC). Following elution, chromatin was reverse-crosslinked with Proteinase K (20 mg/mL, ThermoFischer Scientific, Cat# 25530049) and 0.1 mg/mL RNAse A (Sigma-Aldrich, Cat# 10109169001) and DNA was purified by QIAquick PCR kit (QIAGEN). The AsiSI cutting sites on Chromosome 1, 6 and 22 were amplified by qPCR (Primers) and were previously described(Iacovoni *et al*., 2010). The following antibodies were used: anti-H2AK119ub and rabbit IgG Isotype as control (see key sources table).

### DNA end resection by native IdU staining

MP41 cells were seeded on Poly-D-Lysine coated coverslips and transfected as described above. 24 hours before X-ray (5 Gy), cells were labelled with 25 µM IdU (5-Iodo-2’-deoxyuridine). Following X-Ray irradiation, cells were washed with cold 1x PBS, and cytoplasm was pre-extracted with cytoskeleton buffer (CSK) (10 mM PIPES, 100 mM NaCl, 3 mM MgCl2, 300 mM Sucrose, 0.5% Triton-X-100) for 5 minutes on ice. The nuclei were then fixed with Paraformaldehyde (PFA) 4% in PBS for 10 min, washed twice with 1x PBS and blocked with 5% Bovine Serum Albumin (BSA) in PBS for 2 hours at RT. Primary antibodies were incubated overnight at 4°C (mouse-anti-BrdU,BD Biosciences, Cat# 347580; rabbit-anti-Cyclin A2, Cell Signaling Technology, Cat# 67955). The slides were then washed 3 times with 1x PBS and incubate with the appropriate Alexa Fluor secondary antibody and DAPI to counterstain the nuclei. Slides were acquired as described in the immunofluorescence section.

### Immunofluorescence

MP41 and SP6.5 cells were seeded on circular glass slides and treated with bleomycin/phleomycin or irradiated as reported before. Samples were fixed in Paraformaldehyde (PFA) 4% in PBS (phosphate-buffered saline) for 15 minutes at RT and permeabilized with 0.01% Triton™ X-100 (Sigma, Cat# T8787) for 10 minutes at RT. Blocking was performed with 2% Bovine Serum Albumin (BSA) in PBS 1x for 2h at RT and slides were then incubated with primary antibodies for detecting γH2AX, RPA, RAD51, 53BP1 and cyclinA2, overnight at 4°C (see key resources table). After 3 washes with PBS 1x, slides were incubated with anti-mouse or anti-rabbit Alexa Fluor secondary antibody (Invitrogen) and DAPI to counterstain nuclei for 1h at RT. Following 3 washes with PBS 1x, samples were mounted with ProLong™ Gold Antifade mountant (Invitrogen, Cat# P36934) and acquired using z-stack mode with a point laser scanning confocal microscope (Zeiss LSM 700, ZEN software). All the acquired images were processed using FIJI software (ImageJ).

### Bioinformatic analyses

Data concerning patients were retrieved from QIMR Oncotarget 2016(Johansson *et al*, 2016), TCGA Firehose Legacy, TCGA GDC 2025(Nguyen *et al*, 2022), MSK Cell 2021, PanCancer Atlas(Ellrott *et al*, 2018; Hoadley *et al*, 2018; Taylor *et al*, 2018), MSK Hepatology 2021 (intrahepatic cholangiocarcinoma)(Boerner *et al*, 2021) by cBioportal database (http://www.cbioportal.org). Patient cohort for all the analyzed datasets was stratified according to BAP1 status (diploid WT, -/Mut and -/- or WT vs mut).

### Quantification and statistical analyses

All graphs and statistical analyses were performed using GraphPad Prism 10 (GraphPad Software Inc.). All results are expressed as median or mean (where indicated). Statistical analysis was performed by using unpaired or paired t-test, Ordinary one-way ANOVA and Two-way Anova. For Kaplan–Meier analysis, Mantel-Cox test was applied. Except where noted, all experiments involved three biological replicates.

---

**Supplementary Figure S1. Impact of BAP1 mutation in cancer**.

(A) Overall survival curve in BAP1 +/+ and BAP1 -/mut UM patients stratified by BAP1 mutations in functional (catalytic domain, from 1 to 200 aminoacids) vs non-functional domains. Data were retrieved from TCGA GDC 2025 in cBioportal database.

(B) Kaplan–Meier survival curve for uveal melanoma patients with BAP1 wild-type (green) and mutated BAP1 (red) tumors. P-value is indicated. Cohorts retrieved from MSK Cell 2021 in cBioportal database(Nguyen *et al*., 2022).

(C) Kaplan–Meier survival curve for mesothelioma patients with diploid BAP1 wild-type (green), BAP1 null/mutated (red) and BAP1-null (blue) tumors. P-values are indicated. Cohorts retrieved from TCGA PanCancer Atlas.

(D-G) Violin plots showing aneuploidy score, fraction of genome altered, microsatellite instability (MSI) score and mutation count for BAP1 +/+ versus BAP1-null/mutated or BAP1-null tumors in mesothelioma (D, E), ICC (F), and ccRCC (G). Red line: median. ****p < 0.0001, *\*p < 0*.*05*, ns: not significant. P-values by Ordinary One-way Anova (D, E,G) and Unpaired t-test (F). Cohorts retrieved from TCGA PanCancer Atlas (mesothelioma and ccRCC) and MSK Hepatology 2021 (ICC) from cBioportal.

**Supplementary Figure S2. BAP1 loss impacts on the resolution kinetics of DSBs in MP41 cells**.

(A,B) Western blot showing yH2AX levels during recovery from radiomimetic-induced DSB in MP41(A) and SP6.5(B).

(C,D) Western blots confirming BAP1-depletion in MP41 (C) and SP6.5 (D) cells stained for yH2AX.

**Supplementary Figure S3. BAP1 depletion impairs 53BP1 recruitment after DSB induction in SP6.5 cells**.

(A) Western blot confirming BAP1 and BRCA1 knockdown at 72 h post-siRNA transfection in MP41 cells stained for RAD51/53BP1.

(B, E) Representative immunofluorescence images of 53BP1 (green) in SP6.5 cells following BAP1 depletion at the indicated hours during recovery from Phleomycin treatment (B) or post-IR (5 Gy) (E). DAPI stains DNA. Scale bar: 10 µm.

(C, F) Scatter dot plots showing the number of 53BP1 (F) foci per cell (shown as dots) in SP6.5 cells following phleomycin treatment (C) or IR (F) across two independent experiments. Red line: mean. ****p < 0.0001. P-values by Ordinary one-way ANOVA.

(D, G) Violin plots showing the percentage of 53BP1+ cells (>5 foci/nucleus) over the total population per field (dots) during recovery from Phleomycin treatment (D) or post-IR (G) in SP6.5 cells. Two independent experiments were performed. Red line: median. ****p < 0.0001, **p < 0.01, *p < 0.05. P-values by one-Ordinary way ANOVA.

**Supplementary Figure S4. BAP1 depletion compromises HR and NHEJ repair efficiency**.

(A) Western blot confirming BAP1 and BRCA1 knockdown at 72 h post-siRNA transfection in U2OS HR-GFP/5′EGFP cells.

(B,D) Flow cytometry plots showing RFP+ populations (yellow gate) following I-SceI-RFP transfection in HR-GFP/5’EGFP U2OS (B) or PIMEJ5GFP U2OS (D) clones.

(C) Representative qPCR bar plot confirming BAP1 knockdown in U2OS PIMEJ5GFP cells at 72 h post-siRNA transfection.

**Supplementary Figure S5. BAP1 depletion impairs HR and SSA repair efficiency in HR-GFP/3**′**EGFP U2OS cells**.

(A) Schematic of the integrated HR-EGFP/3′EGFP reporter construct in the selected U2OS clone.

(B) Western blot confirming BAP1 and BRCA1 knockdown at 72 h post-siRNA transfection in U2OS HR-EGFP/3′EGFP cells.

(C) Bar plot showing HR/SSA repair efficiency across 3 independent experiments. Data normalized to control and shown as mean ± SD. ****p < 0.0001. P-values by Ordinary one-way Anova.

(D) Representative flow cytometry plots showing the percentage of RFP+ cells (yellow gate, left panels) transfected with I-SceI-RFP within the total population, and GFP+ cells within the RFP+ gate (right panels). RFP: Y585-PE-A; GFP: B525-FITC-A, SSC: side scatter.

**Supplementary Figure S6. BAP1 depletion leads to aberrant RPA recruitment following DSB in SP6.5 cells**

(A) Cell cycle analysis on synchronised MP41 showing the fraction of G1, S and G2 cells. Cells were collected at 6 hours post Nocodazole release.

(B) Western blot showing the H2AK15ub levels in MP41 following IR (10 Gy) (left panel) and H2BK120ub levels upon BAP1 depletion in G1 cells. MP41 were irradiated (5 Gy) at 6 hours from Nocodazole release and collected 3 hours post IR (right panel).

(C) Real time PCR plot confirming BAP1 knockdown in DIvA cells in 2 independent experiments.

(D) Scatter dot plots showing the number of RPA foci per cell during the indicated recovery time points from bleomycin treatment in MP41. Red line: Mean, ***p < 0.001, **p < 0.01, p-Value by Ordinary One-way Anova. (left panel). Representative images for RPA immunofluorescence staining (right panel). DAPI stains DNA. Scale bar: 10 µm.

(E, F) Scatter dot plots showing the number of RPA+ CyclinA2 ^−^ cells over the total fraction of cyclinA2^-^ cells per field (left) and the quantification of RPA foci in cyclin A2^-^ cells (middle) in SP6.5 following Phleomycin treatment or IR. Red line: Mean. **p < 0.01, *p < 0.05. P-values by Ordinary one-way Anova. Representative immunofluorescence images for CyclinA2 (red), RPA (green) co-staining in SP6.5 cells (right). Arrows mark Cyclin A2^−^ (G1 phase) cells with RPA foci. DAPI stains nuclei, Scale bar: 10 µm.

**Supplementary Figure S7. PTC-209 treatment modulates DSB repair defects in BAP1-depleted cells**.

(A) Real time PCR confirming BAP1 knockdown in MP41 cells for non-denaturing IdU assay. Two independent experiments are shown.

(B) qPCR bar plots showing BAP1 and ctIP knockdown in MP41 cells. Data shown as mean + SD of technical replicates. 2 independent experiments were conducted.

(C) Representative immunofluorescence images showing AsiSI-induced DSB via yH2AX foci detection (red) after 4 hours following the addition of 300 nM 4-OHT. DAPI stains nuclei. Scale bar 10 µm.

(D, E) Western blot confirming BAP1 knockdown in DivA (D) and MP41 (E) cells treated with PTC-209 for ChIP-qPCR and Cyclin A2/RPA immunofluorescence staining shown in Figure 5.

**Supplementary Figure S8 and S9. Original full scan of western blots**.

