## Supplementary figures and images for "BAP1 loss impairs Non-Homologous End Joining DNA repair promoting genomic instability"

### Supp. Figures 1-9

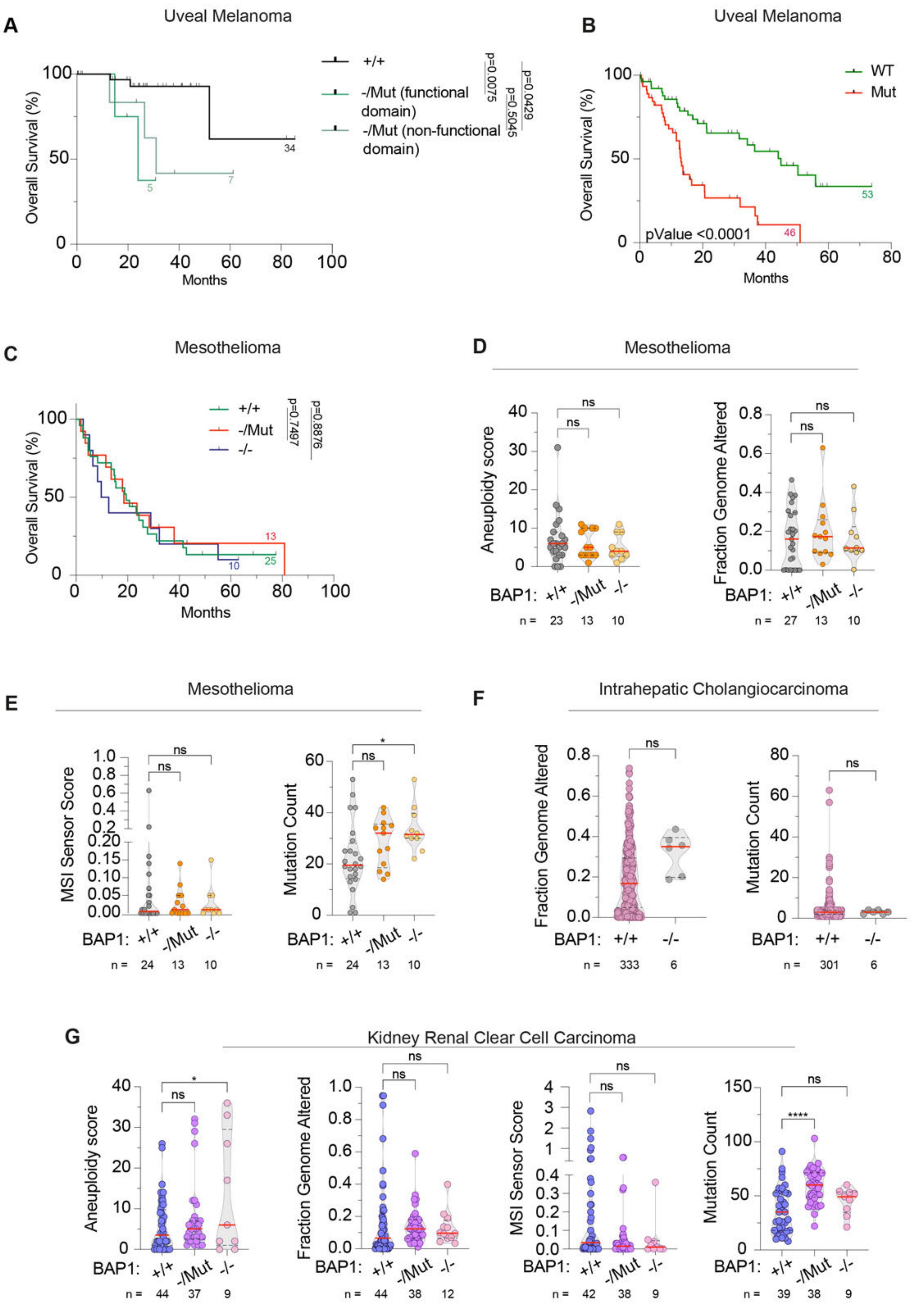

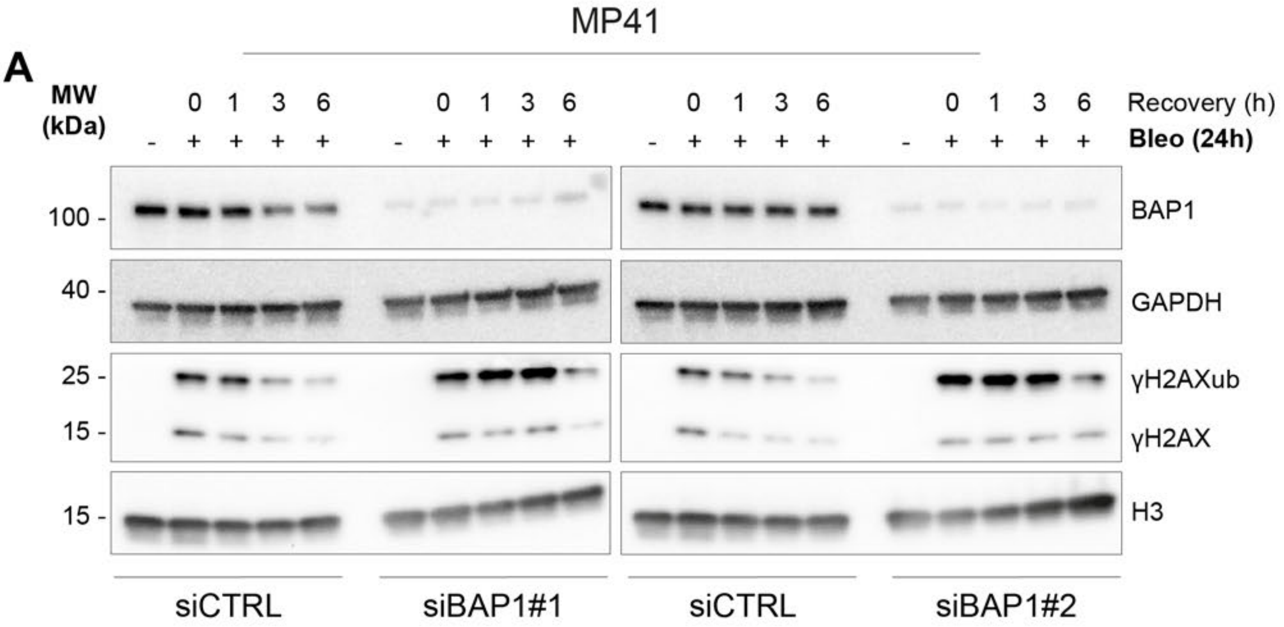

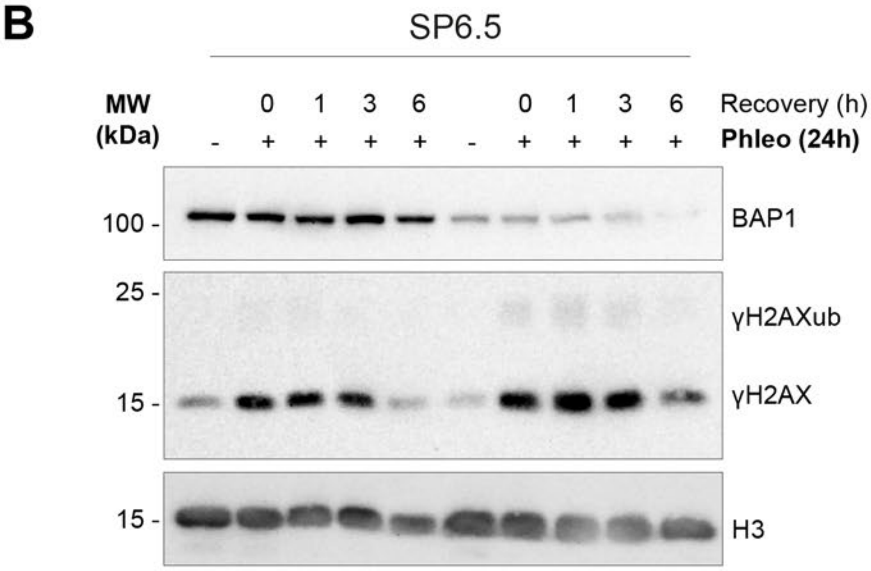

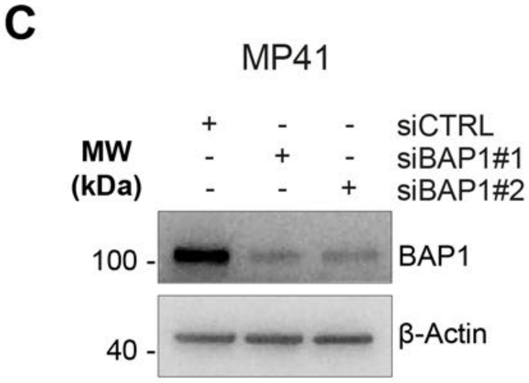

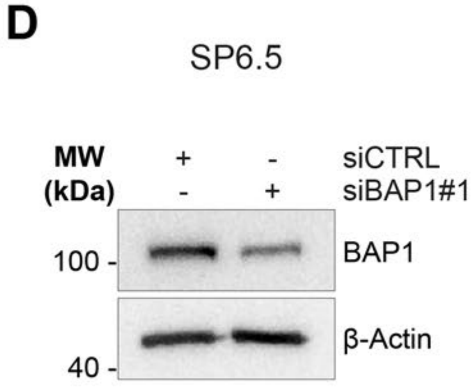

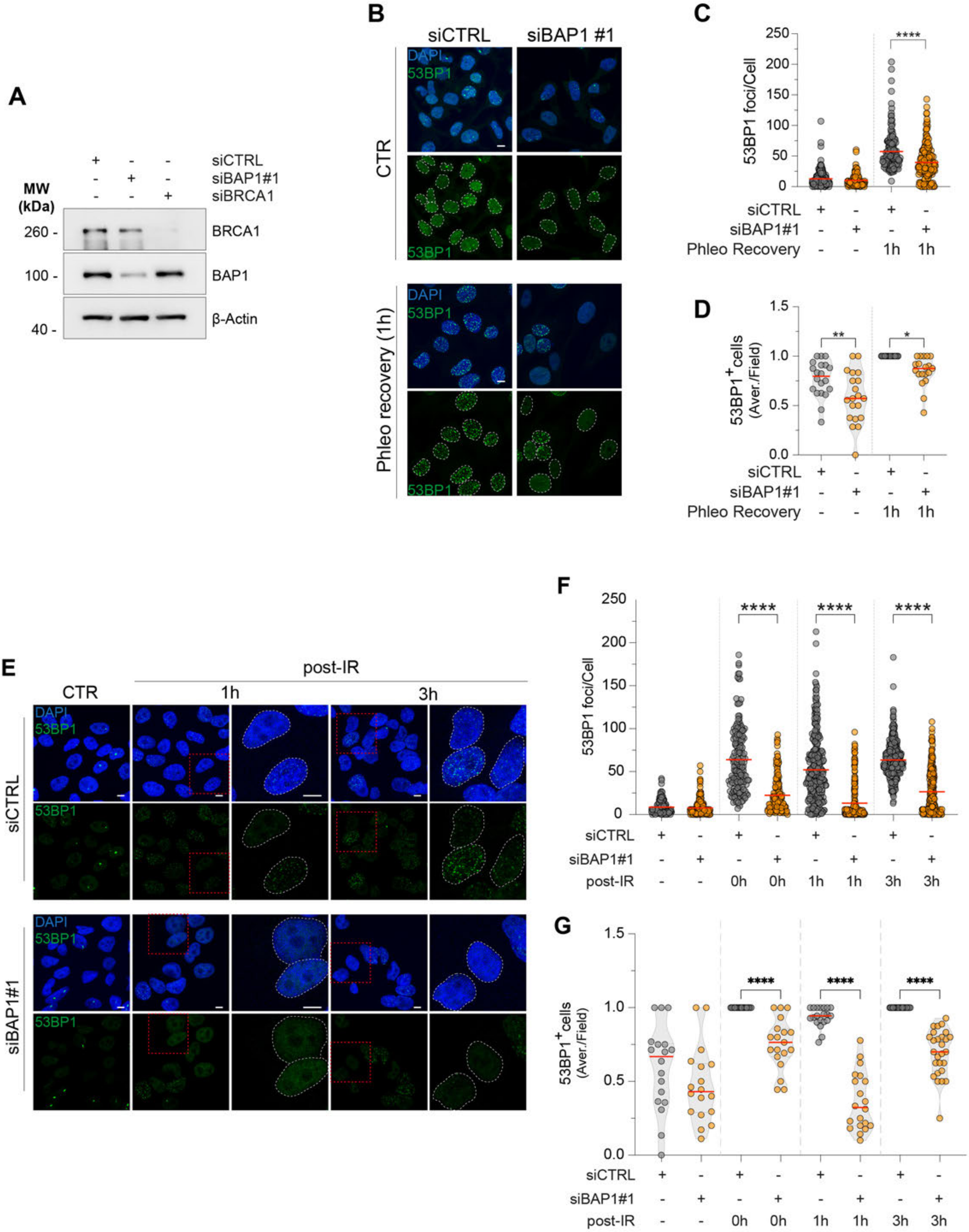

#### FIGURE S4

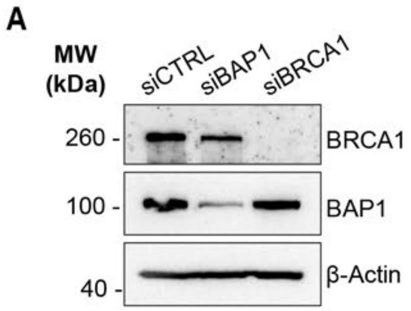


HR-GFP/5'EGFP

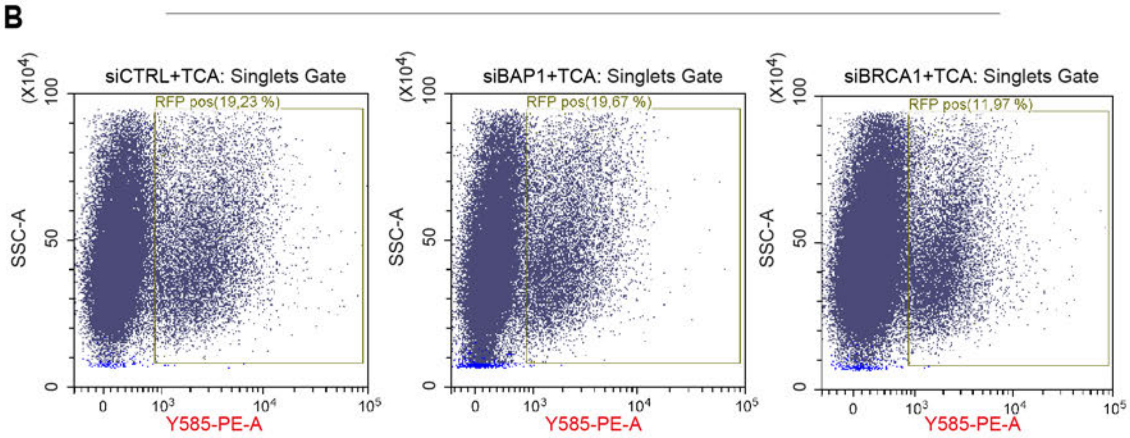


PIMEJ5GFP

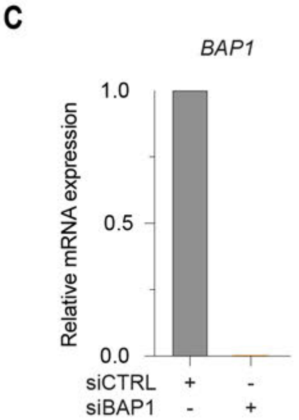


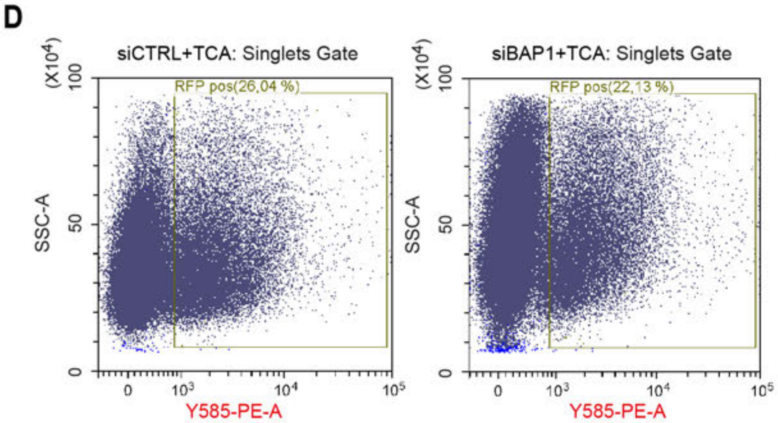

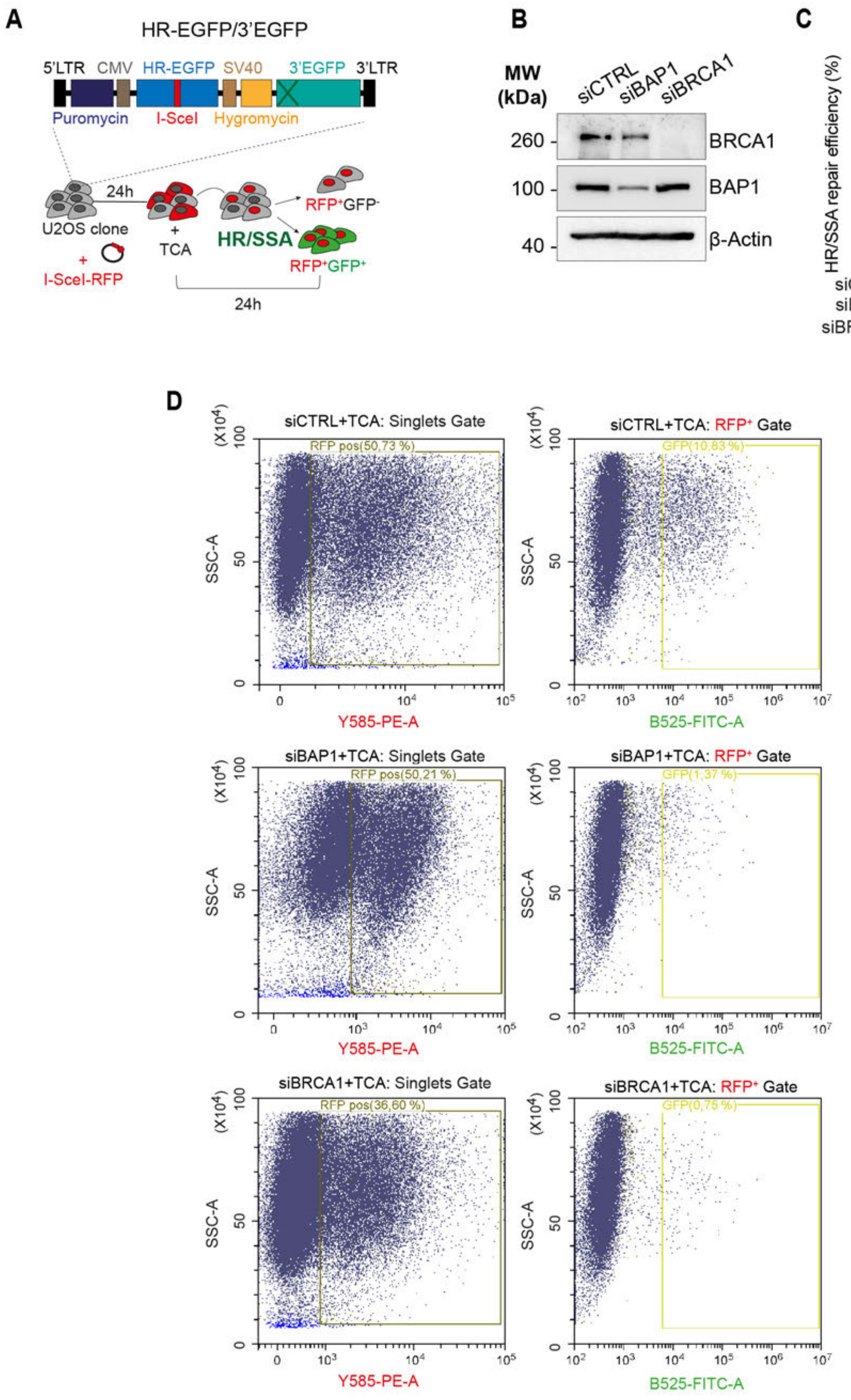

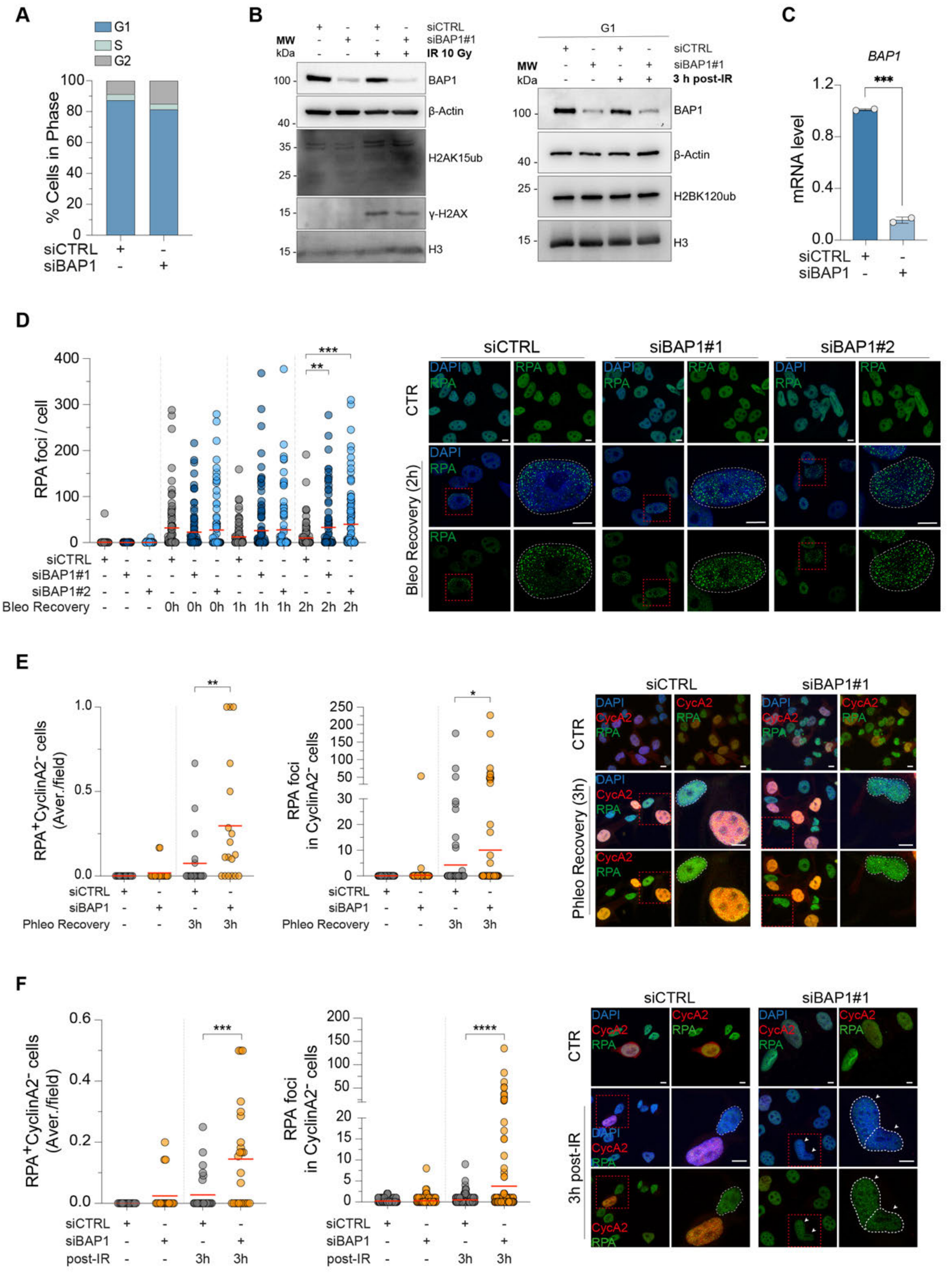

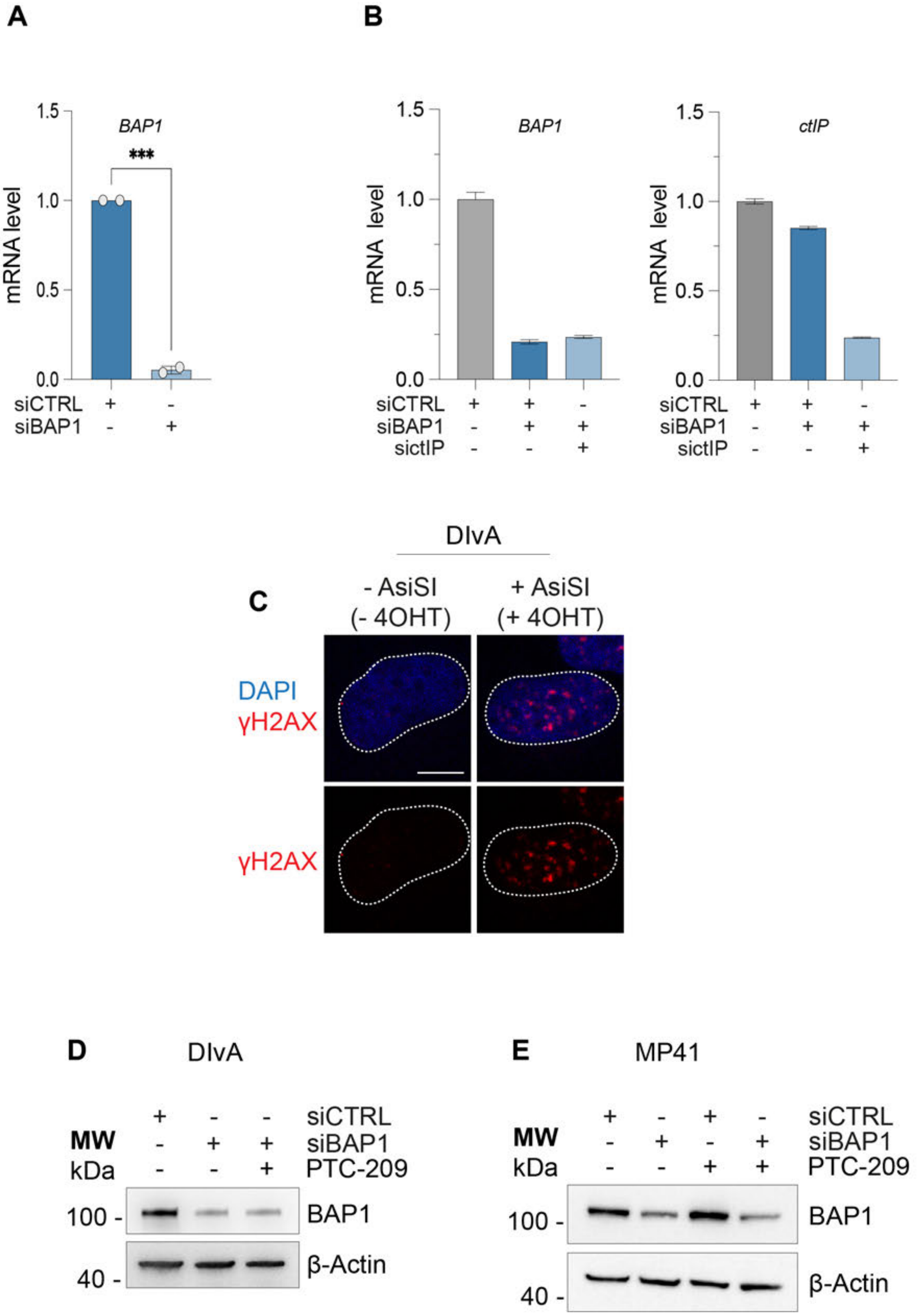

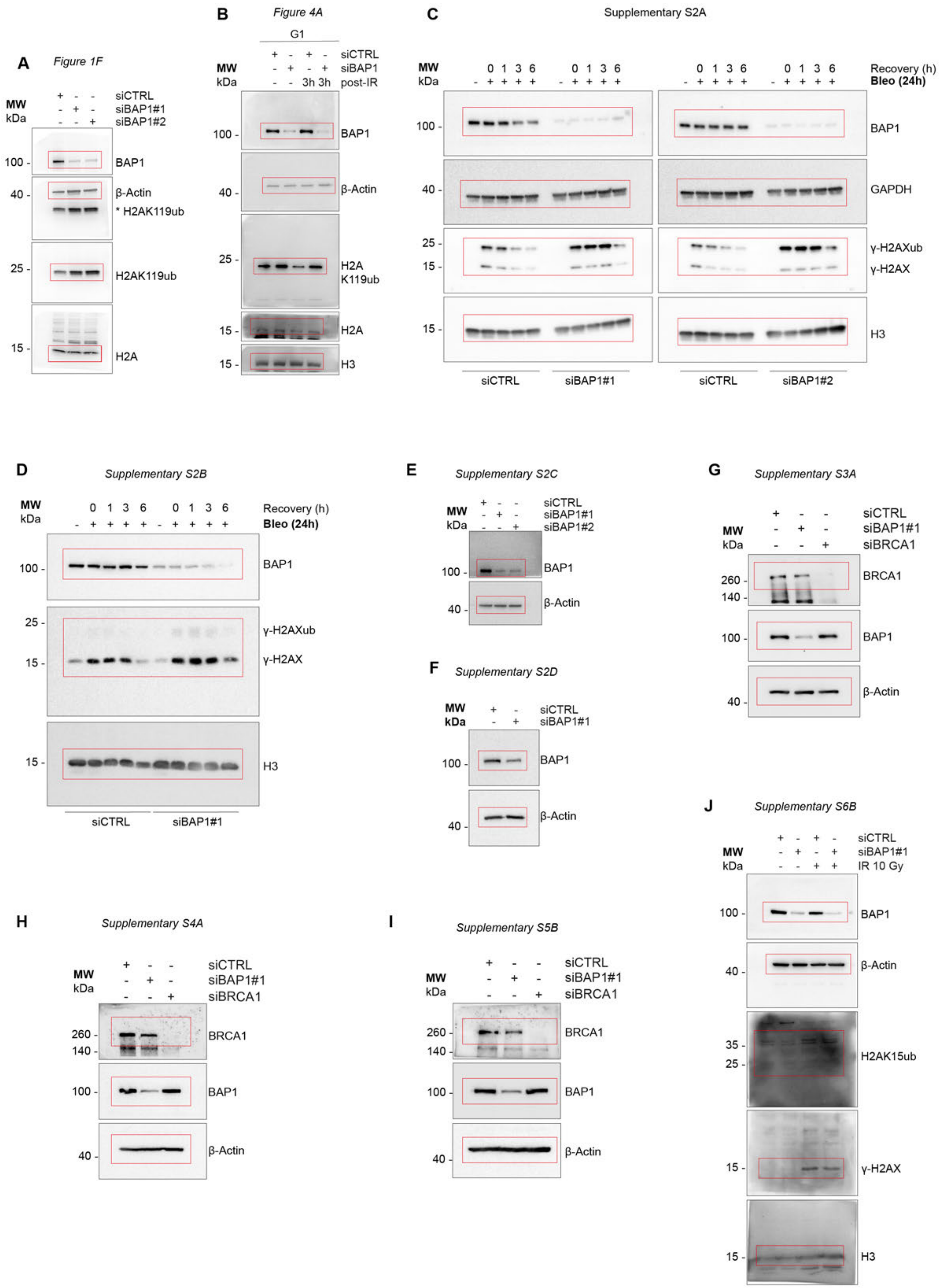

**Figure S9**

*Figure S6B*

**A**

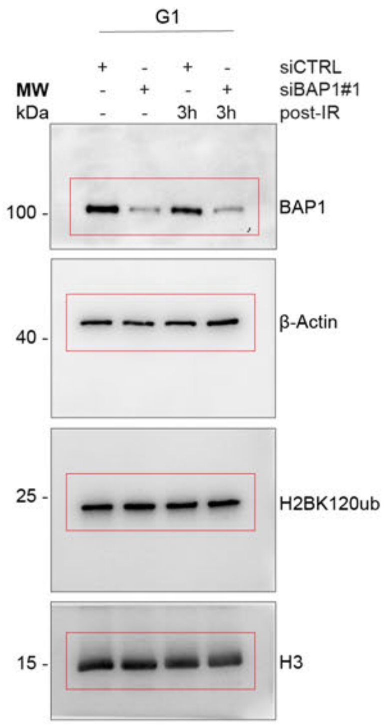


**B** *Figure S7D*

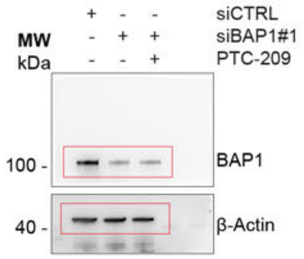


**C** *Figure S7E*

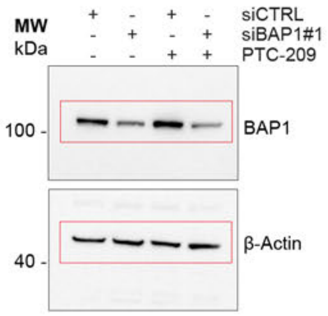
